# Stem-cell niche self-restricts the signaling range via receptor-ligand degradation

**DOI:** 10.1101/414078

**Authors:** Sophia Ladyzhets, Matthew Antel, Taylor Simao, Nathan Gasek, Mayu Inaba

## Abstract

Stem-cell niche signaling is short-range in nature, such that only stem cells but not their differentiating progeny experience self-renewing signals. At the apical tip of the *Drosophila* testes, 8 to 10 germline stem cells (GSCs) surround the hub, a cluster of somatic cells that function as the major component of the stem cell niche. We have shown that GSCs form microtubule-based nanotubes (MT-nanotubes), which project into the hub cells, serving as the platform for niche signal reception: the receptor Tkv expressed by GSCs localizes to the surface of MT-nanotubes, where it receives the hub-derived ligand Decapentaplegic (Dpp), ensuring the reception of the ligand specifically by stem cells but not by differentiating cells.

Here we show that receptor (Tkv)-ligand (Dpp) interaction at the surface of MT-nanotubes serves a second purpose of dampening the niche signaling: we found that the receptor Tkv and the ligand Dpp are internalized into hub cells and are degraded in the hub cell lysosomes. Perturbation of hub lysosomal function or MT-nanotube formation leads to excess receptor retention within germ cells as well as excess Dpp that diffuses out of the hub, leading to ectopic activation of niche signal in differentiating germ cells. Our results demonstrate that MT-nanotubes plays dual roles in ensuring the short-range nature of the niche signaling by 1) providing exclusive interphase of the niche ligand-receptor interaction and 2) limiting the amount of available ligand-receptor via their degradation.

## Introduction

Many stem cells reside in a special microenvironment, called niche, to maintain their identity^1^. Niche signaling is believed to be short-range such that only stem cells but not differentiating cells activate self-renewal signaling above the threshold. In the *Drosophila* testis, germline stem cells (GSCs) reside in the niche formed by post-mitotic somatic cells called hub cells. GSCs typically divide asymmetrically, giving rise to one daughter cell that retains the attachment to the hub to self-renew and the other daughter cell, gonialblast (GB), that is displaced away from the hub and differentiate into spermatogonia (SG). Hub cells secrete ligands, Dpp and Unpaired (Upd). Dpp activates Bone Morphogenetic Protein (BMP) pathway, whereas Upd activates JAK/STAT pathway in GSCs, both of which are required for maintenance of GSCs^2 3 4 5^. These niche-derived ligands must act over a short range so that signaling is only active in GSCs, but not in GBs. Defining the boundary of niche signaling between abutting GSCs and GBs is of critical importance in maintaining stem cell population while ensuring differentiation of their progeny^2^. However, how this short-range nature of niche signaling is achieved is poorly understood.

We have previously demonstrated that the cellular projections, microtubule-based (MT) - nanotubes present on GSCs project into hub cells (Figure 1A). Tkv is produced by GSCs and trafficked to the surface of MT-nanotubes where it interacts with hub-derived Dpp, leading us to propose that MT-nanotubes serves as a signaling platform for productive Dpp-Tkv interaction (Figure 1A). Because MT-nanotubes are formed only by GSCs, this likely contributes to short-range nature of the niche signaling by excluding GBs from receiving Dpp. Here we show that MT-nanotubes serve a second function: in addition to serving as a platform for Dpp-Tkv interaction, it also promotes internalization of Tkv-Dpp by hub cells, where they are degraded in lysosomes. Perturbation of hub lysosomes or MT-nanotube formation resulted in signal activation in non-stem cell populations, suggesting that Dpp-Tkv degradation in the hub plays a critical role in dampening the signal outside the niche. We propose here that MT-nanotubes not only promote the specificity of niche signal reception by stem cells, but also limits the range of niche signal by degrading the receptor-ligand complex.

**Figure 1.**
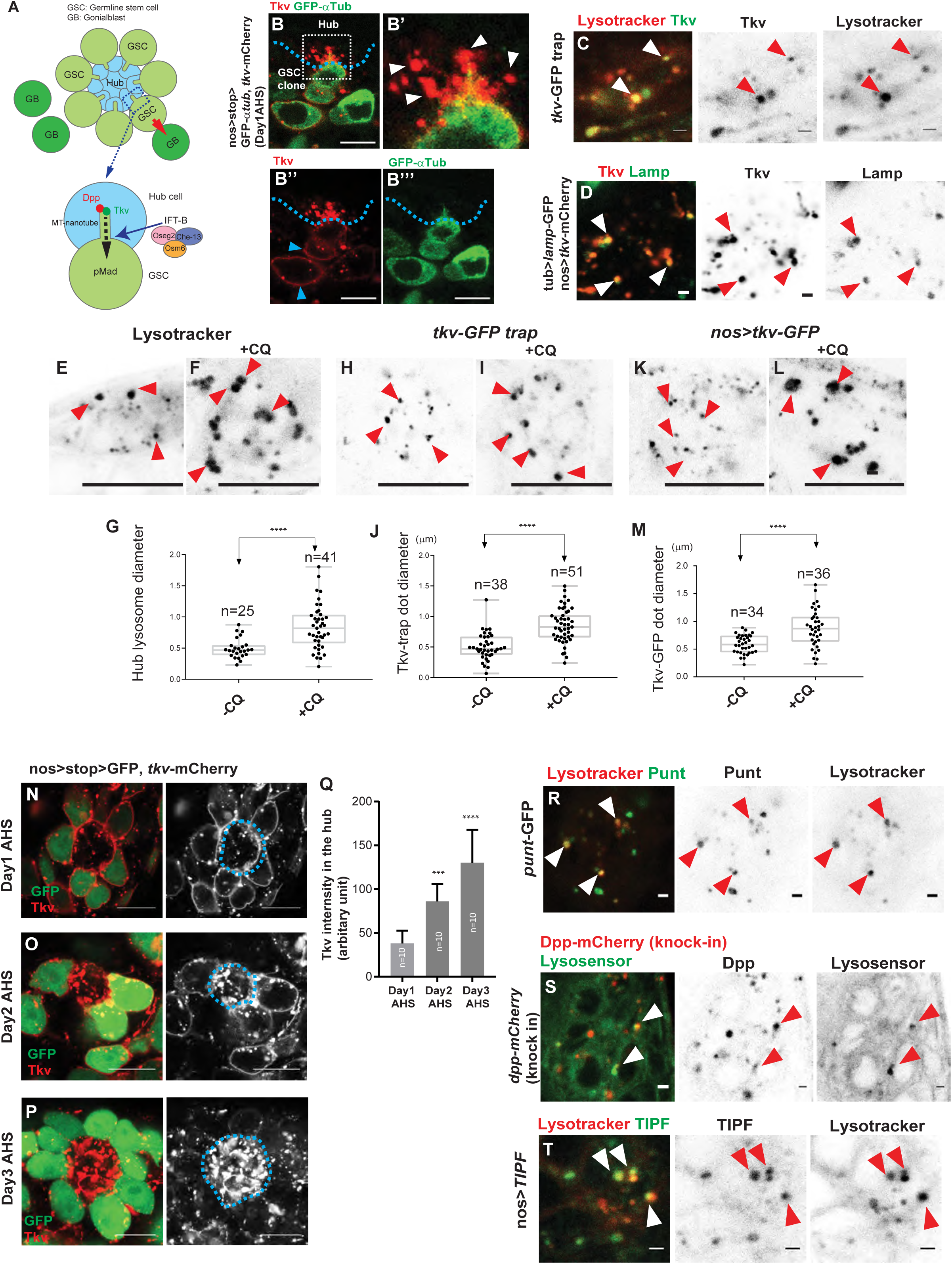
Tkv receptor expressed in GSC is transferred to hub lysosomes. **A, Top**, Schematic of the *Drosophila* male GSC niche. GSCs are attached to the hub. The immediate daughters of GSCs, the GBs, are displaced away from the hub then start differentiation (a red arrow indicates the division orientation). **Bottom**, MT-nanotube-mediated Dpp ligand-Tkv receptor interaction at the hub cell-GSC boundary. IFT-B proteins are responsible for MT-nanotube formation/maintenance. **B**, A GSC clone expressing Tkv-mCherry (red) and GFP-αTub (green); white box is inset **B’**. An arrow in **B”’** indicates an MT-nanotube illuminated by GFP-αTub and Tkv-mCherry. White arrowheads indicate localization of Tkv-mCherry outside of the MT-nanotube within hub cells (**B’**). Blue arrowheads indicate retention of Tkv-mCherry in plasma membrane of differentiating cells (**B’’**). Blue dotted lines outline hub-GSC boundaries. **C, D**, Representative images of hub area of indicated genotypes. Arrowheads indicate the localization of Tkv in the hub lysosomes (Marked by lysotracker in **C**, Lamp-GFP in **D**). **E, F**, Representative images of lysotracker-positive lysosomes in the hub in 4-hour cultured testes without (**E**) or with (**F**) chloroquine (+CQ). **G**, Measured diameters of lysosomes in the hub with or without CQ. **H, I**, Representative images of the hub area of tkv-GFP trap testes in 4-hour cultured testes without (**H**) or with (**I**) chloroquine (+CQ). **J**, Measured diameters of Tkv punctae with or without CQ. **K, L**, Representative images of the hub area surrounded by Tkv-GFP-expressing GSCs under nosGal4 driver in 4-hour cultured testes without (**K**) or with (**L**) chloroquine (+CQ). Red arrowheads indicate Tkv-positive punctae. **M**, Measured diameters of Tkv punctae in the hub with or without CQ. **G, J, M**, largest diameter chosen from 0.5 μm interval z-stacks for each dot. Indicated numbers of punctae from two independent experiments were scored for each group. The indicated numbers of dots from two independent experiments were scored for each data point. Data are means and standard deviations. *P* values were calculated by Student’s t-tests (****P<0.0001) The indicated numbers (n) of dots from two independent experiments were scored for each group. Box plot shows 25–75% (box), median (band inside), and minima to maxima (whiskers). **N-P**, Representative images of hub and surrounding GSCs at the indicated days after induction of Tkv-mCherry expression. Hub is encircled by a blue dotted line. **Q**, Measured intensities of Tkv-mCherry signal in entire hub area (the largest plane was selected for measurement from 0.5 μm interval z-stacks) at the indicated days after induction of Tkv-mCherry expression. The indicated numbers (n) of testes from two independent experiments were scored for each group. Data are means and standard deviations. The adjusted *P* values from Dunnett’s multiple comparisons test are provided as ***P<0.001, ****P<0.0001. **R-T**, Representative images of hub area of indicated genotypes. Arrowheads indicate the localization of Punt (**R**), Dpp (**S**) and TIPF (**T**) in the hub lysosomes (Marked by lysotracker in **R** and **T**, lysosensor in **S**). Scale bars, 10 μm in **B, D I-P** and 1 μm in **G-H, R-T**.

## Results

### Tkv receptors expressed in GSC are transferred to hub lysosomes

We have previously shown that Tkv, the receptor for one of the hub-derived self-renewal ligands, Dpp, is produced in GSCs and trafficked to the surface of the MT-nanotubes. At the surface of MT-nanotubes, Tkv engages in signaling by interacting with Dpp derived from hub cells. We showed that *Intraflagellar transport-B* (*IFT-B*; *oseg2, osm6*, and *che-13*) genes are required for MT-nanotube formation^6^. In the absence/reduction of MT-nanotubes, Tkv is retained in the cell body of GSCs, leading to compromised Dpp-Tkv interaction and thus reduced the activation of downstream BMP signal^6^.

In addition to Tkv localization on MT-nanotube surface, we noticed that Tkv-mCherry expressed in germline were observed within the cell body of hub cells, not necessarily colocalizing with the MT-nanotubes (Figure 1B, white arrowheads). We found that this Tkv population found in the hub cells colocalizes with lysosomes (Figure 1C, D). Tkv’s lysosome localization was further examined by using chloroquine (CQ) treatment, a drug that raises lysosomal pH and inhibits lysosomal enzymes^7–9^. CQ treatment is known to enlarge the lysosome size^10^ (Figure 1E-G): when testes were treated by CQ, we observed that both endogenously tagged Tkv (Tkv-GFP trap) and germline expressed tkv-GFP localized to the enlarged punctae in the hub (Figure 1H-M), confirming that a large proportion of Tkv positive punctae are lysosomes. It should be noted that Tkv-GFP trap, which showed similar lysosomal localization pattern, exhibited complete overlap with germline expression of Tkv-mCherry transgene (Figure S1A), indicating that Tkv-GFP trap signal seen in the hub is all originated in GSCs and Tkv localization in hub lysosomes is unlikely due to the overexpression artifact.

To further confirm the translocation of Tkv from one cell to the other (GSC to hub cells), we followed the temporal order of Tkv localization after its production by inducing GSC clones expressing Tkv-mCherry. When GSC clones expressing Tkv-mCherry was induced by heat shock (hs) (see methods), Tkv-mCherry was first observed on the GSC plasma membrane and in the GSC cytoplasm as puncta (day 1 after hs). After 2 days post hs, its signal in the hub became evident. Finally, after 3 days, the Tkv signal along the GSC plasma membrane reduced and the Tkv signal in the hub increased further (Figure 1N-Q). These results indicate that Tkv is transferred from GSC to the hub cells.

In addition to Tkv, the type II receptor Punt, a co-receptor of Tkv, was observed in the hub lysosomes (Figure 1R). Moreover, Dpp ligand (visualized via a knock-in line in which endogenous Dpp is fused to mCherry^11^ and completely colocalized with Tkv-GFP trap in the hub, Figure S1B), was also seen in lysosomes in the hub (Figure 1S). A reporter of ligand-bound Tkv, TIPF^12^ was also the same (Figure 1T). Taken together, these results suggest that GSC-derived Tkv is trafficked to the MT-nanotubes and then to lysosomes in the hub, together with co-receptor Punt and the hub-derived ligand Dpp. Based on these results, we hypothesized that Tkv, Punt and Dpp may be degraded within the hub cell lysosomes.

### Hub lysosomes degrade Tkv to downregulate Dpp signal in GSCs

To test the possibility that Tkv and Dpp are degraded within the hub cell lysosomes, we examined the effect of impairing lysosome function in the hub cells. To this end, we knocked down genes required for lysosomal function using a hub cell-specific driver. Spinster (Spin) is a putative late-endosomal/lysosomal efflux permease and a known regulator of lysosomal biogenesis^13^. It has been also reported that Spin regulates Dpp signaling through degrading Tkv in *Drosophila* eye discs^14^. Lysosomal-associated membrane protein-1 (Lamp1) is an abundant protein in the lysosomal membrane that is required for lysosomes to fuse with endosomes^15^. Hub-specific knock down of these genes led to increased sizes of Tkv punctae in the hub (Figure 2A, B, and C), likely reflecting defective degradation of Tkv in the hub lysosome. This led to a significant increase in pMad levels in GSCs and their immediate progeny (Figure 2D, E, and F), indicating that hub lysosomes are responsible for niche signal attenuation. In contrast, germ cell-specific knockdown of these two lysosomal genes did not alter pMad level (Figure 2G, H, and F). It should be noted that *dpp* mRNA levels showed no detectible alteration in lysosomal-defective hub cells (Figure S2), indicating that niche signal attenuation by hub cells is not caused by a change in *dpp* transcription level. These results suggest that hub lysosomes are responsible for the degradation of GSC-derived Tkv receptor, which may serve to attenuate the niche signaling to the appropriate level.

**Figure 2.**
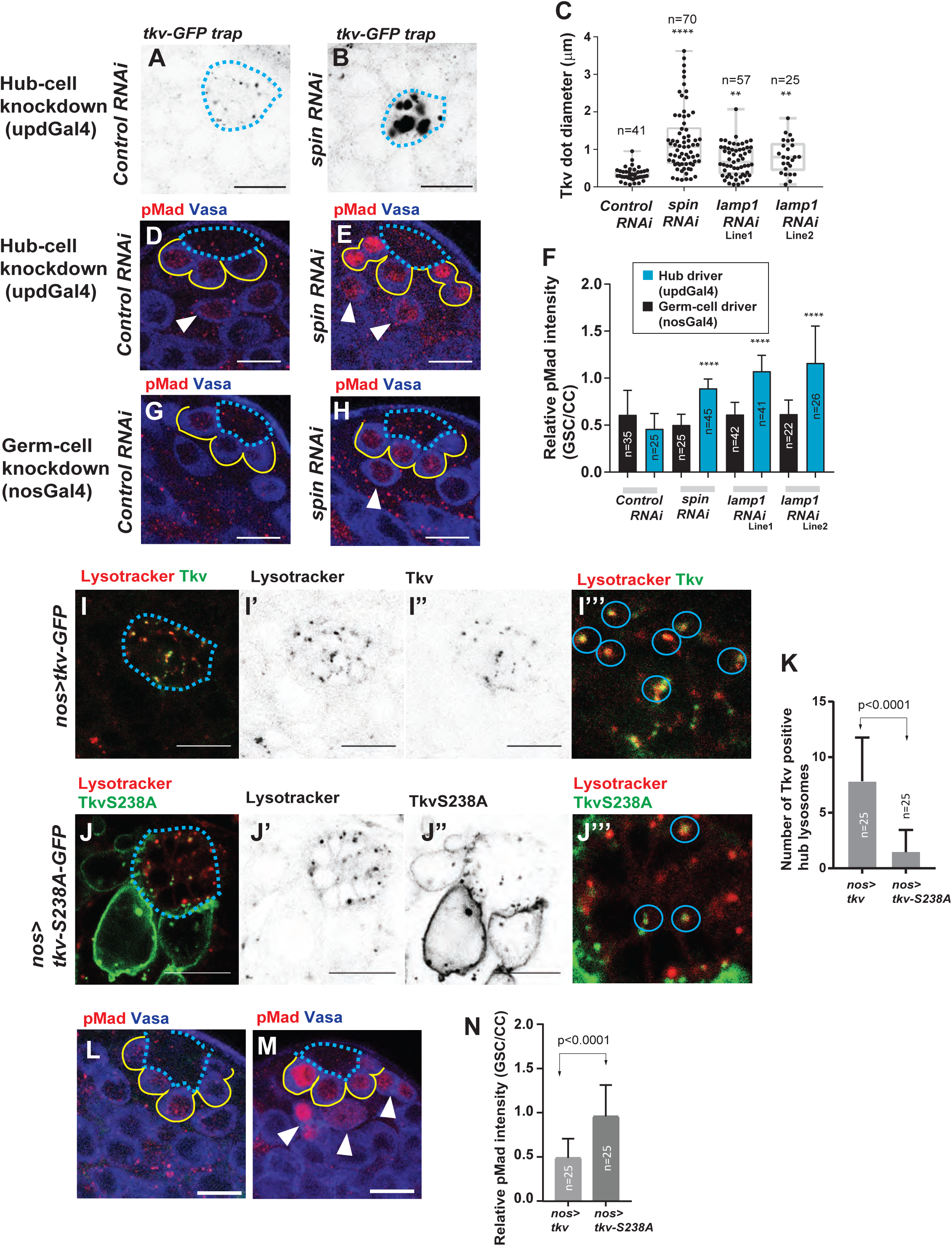
Hub lysosomes degrade Tkv to downregulate Dpp signal in GSC. **A, B**, Representative images of Tkv-GFP trap punctae in the hub (encircled by blue dotted lines) with hub cell-specific RNAi of indicated genes. **C**, Diameters of Tkv-GFP trap punctae in the hub of indicated genotypes (largest diameter for each punctum was chosen by the imaging of 0.5 μm z interval). The indicated numbers (n) of dots from two independent experiments were scored for each group. **D, E**, Representative images of pMad staining (red) in testis of the indicated genotypes. Drivers specific for the hub cells (updGal4) was used to induce expression of the indicated RNAi. Blue dotted lines outline the hub. Yellow lines outline GSCs. White arrowheads in **D, E** indicate germ cells away from the hub that remain pMad-positive. Vasa (blue) is a germ cell marker. **F**, Quantitation of pMad intensity in the GSCs relative to somatic cyst cells (CCs) (Somatic cyst cell is known to be pMad positive, and thus is used for internal control for GSC’s pMad quantification. See Methods). Indicated numbers of GSCs from two independent experiments were scored for each data point. **G, H**, Representative images of pMad staining (red) in testis of the indicated genotypes. Drivers specific for the germ cells (nosGal4) was used to induce expression of the indicated RNAi. Blue dotted lines outline the hub. Yellow lines outline GSCs. Reduction of mRNA level in each RNAi line was Lamp1 Line1; TRiP.HMS01802 47.79%, Line2; TRiP GLV21040 50.87%, Spin 25.05%, see method. **I, J**, Representative images of the testis tips with nosGal4-mediated expression of tkv-GFP or tkvS238A-GFP (**I**, nosGal4>*tkv*-GFP, **J**, nosGal4>*tkv*S238A-GFP). **I”’** and **J’’’** show magnified images of hub area. Lysotracker positive lysosomes (>0.5μm diameter) positive with Tkv are marked by blue circles. **K**, Average number of lysotracker positive lysosomes (>0.5μm diameter) positive with Tkv in the hub counted by the imaging of 0.5 μm z interval throughout entire hub region. **L, M**, Representative images of pMad staining in the testes expressing Tkv-GFP (**L**), or TkvS238A-GFP (**M**). Blue dotted lines outline the hub. Yellow lines outline GSCs. Arrowheads in **M** indicate cells away from the hub which remain pMad (red) positive. **N**, Quantification of pMad intensity in the GSCs (relative to CCs, see Methods) of Tkv- or TkvS238A-expressing testes. The indicated numbers of GSCs from two independent experiments were scored for each group. Scale bars, 10 μm. Data are means and standard deviations. Box plot in **C** shows 25–75% (box), median (band inside), and minima to maxima (whiskers). For **C** and **F**, the P values (adjusted *P* values from Dunnett’s multiple comparisons test) are provided as *P<0.05, **P<0.01, ***P<0.001, ****P<0.0001; or non-significant if not shown (P>0.05). For **K** and **N**, P values were calculated by Student’s t-tests.

### Ubiquitination mediates Tkv degradation by promoting translocation of Tkv from GSC to the hub lysosome

Ubiquitination of membrane proteins is known to be required for recognition by the endosomal sorting complexes required for transport including endocytosis, lysosomal fusion, and degradation^16^. SMAD ubiquitination regulatory factor (Smurf) is a HECT (Homologous to the E6-AP Carboxyl Terminus) domain-containing protein with E3 ubiquitin ligase activity, and disruption of Smurf function has been shown to enhance Dpp-Tkv signal in GSCs^17,18^. It has been reported that phosphorylation of Tkv Ser238 residue is required for Smurf-dependent ubiquitination^17^.

We found that Tkv-S238A-GFP, in which the Serine residue required for Smurf-dependent ubiquitination was mutated, exhibits striking difference from wild type Tkv-GFP on two accounts. First, the protein amount was significantly upregulated in S238A mutant compared to wild type, consistent with its known degradation-resistant nature (Figure2I, J). Moreover, S238A mutant exhibited a localization change: it localized mostly to GSC plasma membrane, and colocalization with the hub lysosome was greatly diminished (Figure2I”’, J”’, K). Consistent with compromised downregulation of Tkv, expression of S238A resulted in upregulation of pMad (Figure2L, M, N).

Taken together, these results demonstrate that ubiquitination of Tkv, induced by phosphorylation of S238, is required for its translocation from GSC to the hub. Our data further suggest that this translocation of Tkv to the lysosome is critical to downregulate excess amount of Tkv.

### Degradation of Dpp-Tkv in the hub lysosome is required for proper differentiation

The above results suggest that degradation of Dpp-Tkv in the hub is required to downregulate Dpp-Tkv signaling. Although, shutting down Dpp-Tkv signal is required for proper differentiation, neither downregulating hub lysosome (*spin* RNAi) nor Tkv trafficking to the hub (IFT-KD), both of which are expected to compromise Dpp-Tkv degradation in the hub, were not enough to impact differentiation. However, we found that compromising Tkv trafficking to the hub (IFT-KD) combined with overexpression of Tkv (TkvOE) (referred to as *IFT-KD/Tkv-OE*) showed prolonged pMad staining in SG populations (Figure 3A-D). Moreover, it frequently caused undifferentiated germ cells to proliferate ectopically outside the niche (hereafter referred to as germline tumor, Figure 3D’ F), reflecting inability to turn down Dpp-Tkv signaling.

**Figure 3.**
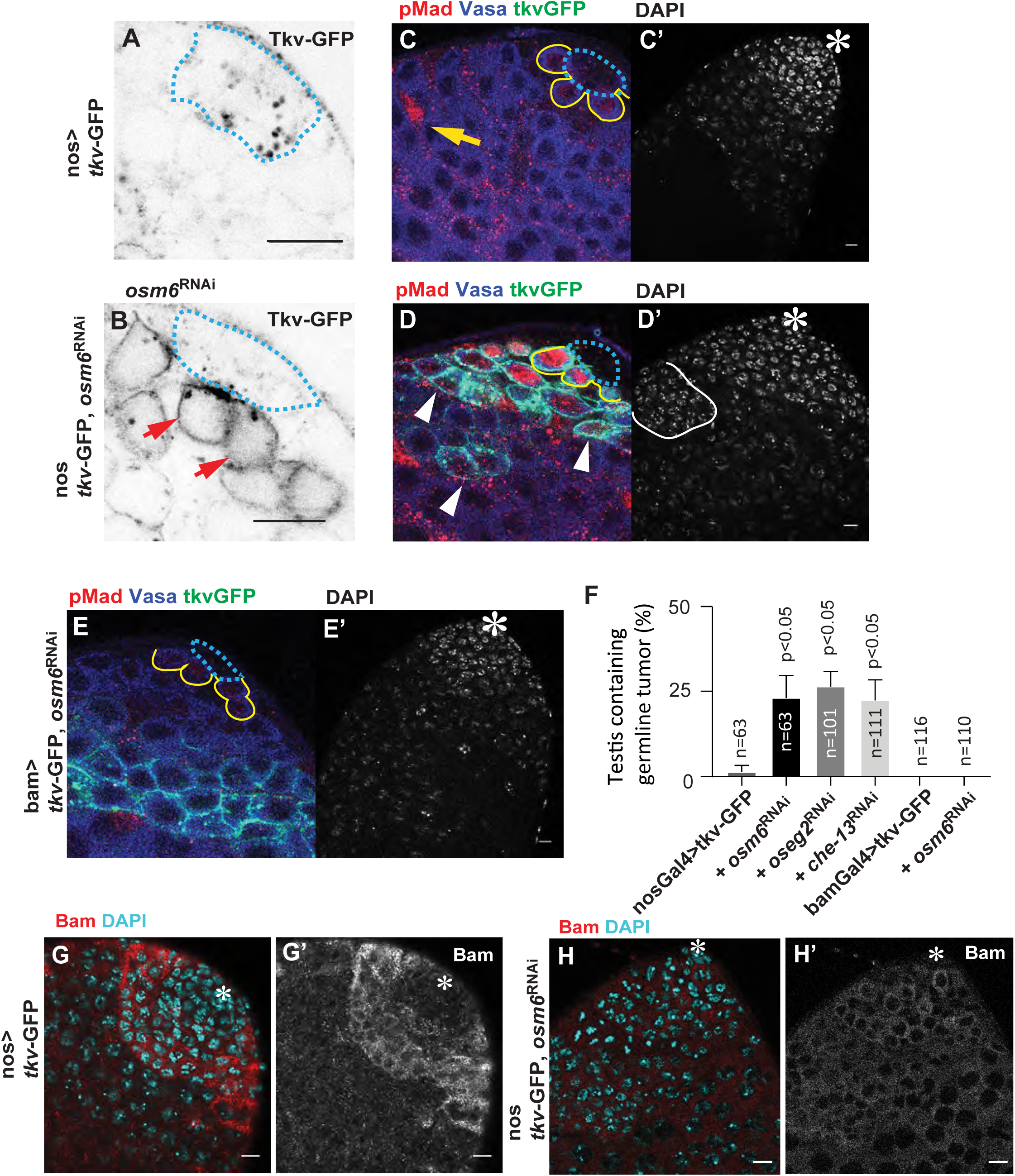
Degradation of Dpp-Tkv in the hub lysosome is required for proper differentiation. **A-D**, Representative images of Tkv-GFP expressing testis tip under the germline specific driver (nosGal4). The hub and surrounding GSCs are outlined by blue dotted lines and yellow lines respectively. **A**, Tkv localizes to punctae in the hub, and almost undetectable level in the GSC cell body. **B**, Tkv-GFP pattern in the tip of testis after knockdown of IFT-B (*osm6* RNAi). Red arrows indicate plasma membrane localization of Tkv. **C, D**, Vasa (blue), pMad (red), Tkv (Green; note that this green signal appears as cyan due to its overlap with Vasa; blue). Note the nosGal4 driven Tkv-GFP seen as punctae in the hub (**A**) is not visible in **C** after the fixation. Yellow arrow in **C** indicates an example of a CC which is known to be pMad positive, and thus is used for internal control for GSC’s pMad quantification. White arrowheads in **D** indicate Tkv-GFP protein retention and prolonged pMad staining in differentiating germ cells. White line in **D’** outlines germline tumor with condensed chromatin detected by DAPI staining, separated from the group of the cells containing GSC, GB, SGs and somatic cyst stem cells which are typically observed near the tip of the testis as a single group (**C’**). **F**, % of testes containing any germline tumor (see **D’**) of the indicated genotypes. The indicated numbers of testes from three independent experiments were scored for each data point. Data are means and standard deviations. Adjusted *P* values from Dunnett’s multiple comparisons test are provided (comparing between control group: nosGal4>tkv-GFP and each group). **G, H**, Representative Bam staining in the (nosGal4>Tkv-GFP) testis without (**G**) or with (**H**) osm6 knockdown. No detectable peak of Bam expression in the testis expressing osm6 RNAi with Tkv-GFP (green). Bam (red); DAPI (cyan). Scale bars, 10 μm. Asterisks indicate the hub.

Although the germline tumors observed in the *IFT-KD/Tkv-OE* testes were located away from the hub (Figure 3D’), tumor formation is likely caused by a defect within the GSC: when expression of *IFT-KD/TKV-OE* was driven in differentiating germline by using the bamGal4 driver (active in 4-8-cell stage SG), it did not lead to tumor formation (Figure 3E, F). These data indicate that GSCs’ inability to shut down Dpp-Tkv signaling due to the expression of *IFT-KD/TKV-OE* is the cause of tumor formation.

Bag of marbles (Bam), a master differentiation factor, whose expression is suppressed by Dpp signal within GSCs, typically peaks around 4-8 SG stage^19^. In the *IFT-KD/Tkv-OE* testes, Bam peak was never observed (Figure 3G, H), consistent with the idea that the germline tumor was caused by ectopic or prolonged Dpp-Tkv signal.

In sum, these results show that combination of compromised Dpp-Tkv degradation and overexpression of Tkv leads to defective attenuation of Dpp-Tkv signaling, revealing the importance of Dpp-Tkv degradation in the hub in promoting differentiation.

### Dpp ligand diffuses out from the hub upon inhibition of lysosomes

If the Tkv digestion in adjacent hub cells is happening only in GSCs, how can IFT knockdown cause a tumor located outside of the niche? We speculate that Dpp ligand may diffuse away from the hub when the lysosomal degradation is impaired. To test this idea, we examined the localization of Dpp-mCherry (mCherry knock-in strain)^11^ after the lysosomal inhibition. CQ treatment increased the size of Dpp-mCherry positive punctae within hub cells (Figure4A, B and E). Likely as a result of defective degradation of Dpp, we detected abundant Dpp-mCherry signal outside the hub after a 4-hour CQ treatment (Figure 4B). Similar results were obtained by using a UAS-Dpp-mCherry transgene expressed specifically in hub cells (Figure 4C, D and F). Moreover, after a 4-hour-CQ treatment, TIPF signal, a reporter of ligand-bound Tkv, was broadly observed on differentiating germ cell membranes located outside of the niche, suggesting the possibility that Dpp diffused out of the niche and bound to Tkv receptor on the cells located far away from the hub (Figure 4G, H). This explains why *IFT-KD/TKV-OE* exhibits germline tumor located away from the hub.

**Figure 4.**
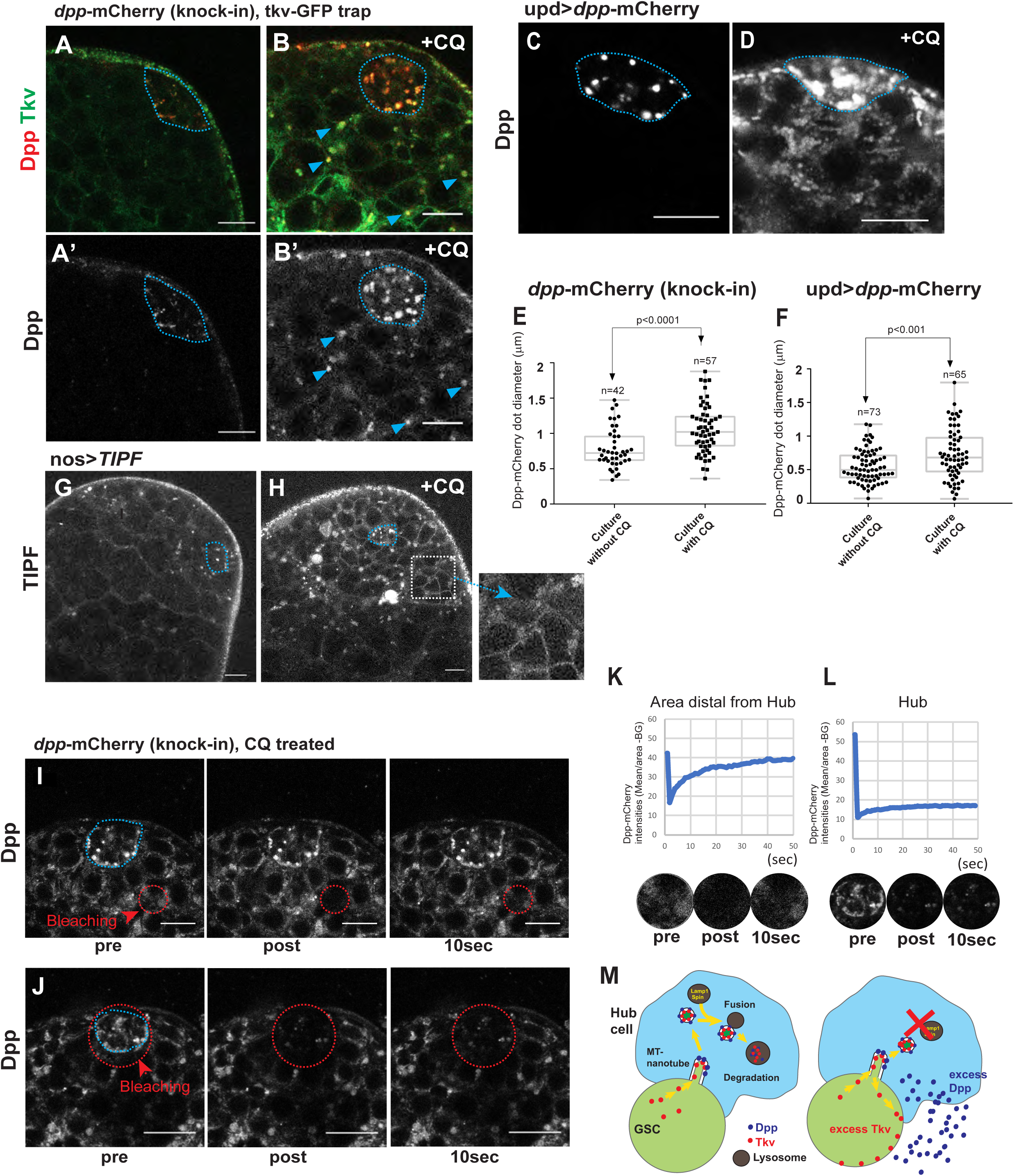
Dpp ligand diffuses out from the hub upon inhibition of lysosomes. **A-D**, Representative images of the testes tips of Dpp-mCherry knock-in with Tkv-GFP trap (**A** and **B**), or the testis expressing Dpp-mCherry under the control of a hub-specific driver (updGal4>*dpp*-mCherry, **C** and **D**) after 4-hour incubation without (**A, C**) or with (**B, D**) CQ (+CQ). Blue arrows indicate lysosomal localization of Dpp in differentiating germ cells. **E, F**, Diameters of Dpp punctae in the hub after incubation with or without CQ. For measurements, only well-isolated dots were chosen to avoid measuring 2 or more overlapped lysosomes. To select single, well-separated lysosomes, lower laser exposure was used if necessary. The largest diameter chosen from 0.5 μm interval z-stacks for each dot. The indicated numbers (n) of dots from two independent experiments were scored for each group. Box plots shows 25–75% (box), median (band inside), and minima to maxima (whiskers). Data are means and standard deviations. *P* values were calculated by Student’s t-test. **G, H**, Representative images of the testis tips with expression of Tkv activation sensor, TIPF, under the control of a germline-specific driver (nosGal4>TIPF) after 4-hour culture without (**G**) or with (**H**) CQ. Insets show magnified images from white square region. **I, J**, Representative FRAP experiments of the testis tips of Dpp-mCherry knock-in after 4-hour chloroquine treatment. The regions encircled by red dotted lines were photobleached and monitored the intensities of mCherry signal before and after the photobleaching at indicated time points. **K, L**, Measured intensities (-BG; background subtracted) of the Dpp-mCherry signal from each region (magnified images from each region are shown under the graph) before and after the photobleaching at indicated time points. The region in **I** represents rapid recovery after bleaching. The region in **J** containing the entire hub area did not show recovery within the full imaging period (50 sec). **M**, Model. **Left;** Tkv expressed in GSCs is transferred to hub lysosomes through MT-nanotubes, where Tkv is subjected to lysosomal degradation. This way, MT-nanotubes contribute to both signal activation and inactivation. **Right;** Hub lysosomal inhibition initiates excess Tkv retention within GSC and excess Dpp ligand starts diffusing out from the niche. Blue dotted lines represent the hub-GSC boundaries. Scale bars, 10 μm.

To determine if the Dpp-mCherry detected outside the hub is due to its diffusion from the hub, we used fluorescence recovery after photobleaching (FRAP) analysis. After photobleaching a 10 μm diameter circle located approximately 10 μm away from the hub center, the photobleached region was quickly equalized by the Dpp-mCherry signal from neighboring regions. The recovery was rapid with an average time of 7.4 ± 2.1 seconds (n=7) to reach 50% of the original intensity (Figure 4I, K, movie1). In contrast, after photobleaching of the entire hub region, signal did not recover (Figure 4J, L, movie2), indicating that the Dpp signal outside the hub is likely the Dpp protein diffused from the hub, and that the hub is likely the sole source of Dpp. It should be noted that the photobleached signal did not always fully recover up to 100% (averaging 70 ± 15%, n=7) possibly because rest of Dpp protein fraction might be trapped in the photobleached field. We speculate that Dpp protein may bind to the extracellular matrix components as reported in *Drosophila* embryo^20^. Alternatively, diffused Dpp ligand might be endocytosed into germ cells after binding to Tkv. Indeed, we observed lysosomal localization of Dpp ligand together with Tkv in germ cells located away from the hub (Figure 4B arrowheads, Figure S3 arrowheads).

In summary, these observations suggest that in the absence of proteolysis of Dpp, the ligand can diffuse from the hub. MT-nanotube-mediated degradation of receptor together with ligand is essential for preventing signal overload outside of the niche.

## Discussion

We have shown previously that niche cells and stem cells interact in a contact-dependent manner, with GSCs and hub cells engaging in productive signaling via MT-nanotubes, enabling highly specific cell-cell interactions and excluding non-stem cells from receiving stem cell signals. Here, we demonstrate MT-nanotubes also contribute to proteolysis of receptor and ligand via transferring the stem cell receptor together with ligand to the niche cell lysosomes (Figure 4M). This ensures the removal of receptor and ligand after the signal engagement and prevents ligand-receptor interactions outside of the niche.

Cytonemes, another type of actin-dependent signaling protrusion^21 22^, also transfer ligand and receptor, allowing the interaction between cells at a distance. Ligand-producing and receptor-producing cells both form cytonemes and both cells have been observed to exchange signaling proteins: receptor into the ligand-producing cells and ligand into the receptor-producing cells^22^. Similarly, exchange of receptor tyrosine kinases, bride of sevenless and sevenless (sev), during *Drosophila* retinal development has also been documented^23^. Exchange of plasma membrane proteins also occurs in mammalian systems; “Trogocytosis” is the phenomena reported in lymphocytes and antigen-presenting cells through the immunological synapse^24 25^. These studies, together with our study, illustrate the universality of such transfer in general contact-dependent signaling.

What could be the potential benefit(s) of this mechanism in which stem cells transfer and degrade their receptor in niche cells instead of digesting by themselves? Since Tkv transport occurs from MT-nanotube membrane, only ligand-bound receptors but not ligand-unbound receptors may be subjected to degradation. In this way, stem cell can secure at least required amount of Tkv. Another possibility is that this might be the strategy of GSCs to avoid receptor endocytosis. Since Tkv is transferred into hub cells, activated receptor never come back to the cell body of GSCs. Such signaling endosomes can act as intracellular platforms for signaling in many other systems^26 27^. BMP signal is also known to utilize signaling endosomes. The activated receptor complex phosphorylates Mad (or receptor-regulated Smad 2 and 3 in mammalian systems) at the C-terminal of BMP receptor presented on the surface of SARA endosomes^28^. Although signaling endosomes can enhance the activation of downstream signal, endosomes can be inherited into differentiating daughter cells after division, which may compromise the specificity of the niche-stem cell signaling. Therefore, transferring the receptor to other cells may be the stem cell-specific strategy to achieve high specificity of the niche signal, such that signal activation only occurs in stem cells, but not in differentiating daughter cells.

It remains to be investigated whether lysosomal proteolysis of ligand and receptor in niche cells, as demonstrated by our study, might also regulate other stem cell systems.

## Acknowledgements

We thank Yukiko Yamashita, Thomas B. Kornberg, Helmut Krämer, Elizabeth R. Gavis, Saugata Roy, Cheng-Yu Lee, Michael Buszczak, the Bloomington Drosophila Stock Center, the Developmental Studies Hybridoma Bank, and the Vienna Drosophila Resource Center for reagents; Yukiko Yamashita, Laurinda Jaffe, Mark Terasaki, Michael Buszczak for discussion; Christopher Bonin for manuscript editing; this work was supported by an NIH grant 1R35GM128678-01 and start-up funds from UConn Health (to M.I.).

## Author Contributions

M.I, Conception and design, acquisition of data, analysis and interpretation of data, drafting and revising the article; S.L, T.S, M.A and N.G., Acquisition of data, analysis and interpretation of data, drafting or revising the manuscript.

## Declaration of Interests

The authors declare no competing interests.

## Methods

### Fly husbandry and strains

All fly stocks were raised on standard Bloomington medium at 25°C (unless temperature control was required), and young flies (0- to 4-day-old adults) were used for all experiments. The following fly stocks were used: tkv-GFP protein trap line (CPTI-002487, inserted in the 1^st^ intron of Tkv-RD) was a gift from B. McCabe. hs-flp; nos-FRTstop-FRT-gal4, UAS–GFP^29^; nosGal4^30^, updGal4 (FBti0002638) were gifts from Y. Yamashita. tubGal80^ts 31^ (gift from C.Y. Lee), bamGal4VP16 (gift from M. Buszczak), tub-GFP-Lamp1^32^ (FBrf0207605, gift from H. Krämer). UAS-TIPF^12^, UAS-Dpp-mCherry ^33^, UAS-Tkv-mCherry ^33^, UAS-Tkv-GFP ^33^ Dpp-mCherry (CRISPR knock-in)^11^ were gifts from T. Kornberg. Stocks from Vienna Drosophila Resource Center (VDRC): Oseg2 RNAi (VDRC GD8122), Osm6 RNAi (VDRC GD24068), Che-13 RNAi (VDRC GD5096), and Punt-GFP (VDRC 318264, 2XTY1-SGFP-V5-preTEV-BLRP-3XFLAG). Other stocks were from Bloomington Stock Center: UAS–GFP–αTubulin (BDSC7253), Spin RNAi (TRiP.JF02782), and Lamp1 RNAi (TRiP.HMS01802 and TRiP GLV21040). For the overexpression of Dpp-mCherry, updGal4^ts^ driver was used, and a combination of updGal4 and tubGal80^ts 31^ was used to avoid lethality. Temperature shift crosses were performed by culturing flies at 18°C to avoid lethality during development and shifted to 29°C upon eclosion for 4 days before analysis. Control crosses for RNAi screening were designed with matching gal4 and UAS copy number using TRiP control stock (BDSC35785) at 25 °C. RNAi screening of candidate genes for Tkv trafficking and degradation was performed by driving UAS-RNAi constructs under the control of nosGal4 or updGal4 (see below for validation method).

### Quantitative reverse transcription PCR

Males carrying nos-gal4 driver were crossed with males of indicated RNAi lines or UAS–GFP–αTubulin (BDSC7253), UAS-Tkv-GFP, or UASp-TkvS238A-GFP transgenic lines. Testes from 100-200 male progenies, aged 0-7 days, were collected and homogenized by pipetting in TRIzol Reagent (ThermoFisher) and RNA was extracted following the manufacturer’s instructions. 1 μg of total RNA was reverse transcribed to cDNA using SuperScript III First-Strand Synthesis Super Mix (ThermoFisher) with Oligo (dT)20 Primer. Quantitative PCR was performed, in duplicate, using SYBR green Applied Biosystems Gene Expression Master Mix on a CFX96 Real-Time PCR Detection System (Bio-Rad). Control primer for αTub84B (5’-TCAGACCTCGAAATCGTAGC-3’/5’-AGCAGTAGAGCTCCCAGCAG-3’) and experimental primer for Spin (5’-GCGAATTTCCAACCGAAAGAG-3’/5’-CGGTTGGTAGGATTGCTTCT-3’), Lamp1 (5’-AACCATATCCGCAACCATCC-3’/5’-CCTCCCTAGCCTCATAGGTAAA-3’), and GFP (5’-GAACCGCATCGAGCTGAA-3’/5’-TGCTTGTCGGCCATGATATAG-3’) were used. Relative quantification was performed using the comparative CT method (ABI manual). Other RNAi lines were validated previously^6^. GFP mRNA levels of nos>Tkv-GFP and nos>TkvS238A were 84.6% and 74.2% of nos>UAS–GFP–αTubulin expression level respectively.

### Generation of pUASP-Tkv S238A transgenic flies

EGFP cDNA was amplified from Drosophila gateway pPGW vector (https://emb.carnegiescience.edu/drosophila-gateway-vector-collection#_Copyright,_Carnegie) using the following primers with restriction sites (underlined): BglII GFP F 5’-AC<u>AGATCT</u>ATGGTGAGCAAGGGCGAGGAGCTGTTCA-3’ AscI GFP R 5’-TA<u>GGCGCGCC</u>TTACTTGTACAGCTCGTCCATGCCGAGA-3’ Products were then digested with BglII and AscI. NotI BglII sites (underlined) were attached to gBlock TkvS238A fragment (Integrated DNA Technologies, sequence as follows). 5’-AT<u>GCGGCCGC</u>ACCATGGCGCCGAAATCCAGAAAGAAGAAGGCTCATGCCCGCTCCCTAACC TGCTACTGCGATGGCAGTTGTCCGGACAATGTAAGCAATGGAACCTGCGAGACCAGACCCGGTGGCAGTTGCTTCAGCGCAGTCCAACAGCTTTACGATGAGACGACCGGGATGTACGAGGAGGAGCGTACATATGGATGCATGCCTCCCGAAGACAACGGTGGTTTTCTCATGTGCAAGGTAGCCGCTGTACCCCACCTGCATGGCAAGAACATTGTCTGCTGCGACAAGGAGGACTTCTGCAACCGTGACCTGTACCCCACCTACACACCCAAGCTGACCACACCAGCGCCGGATTTGCCCGTGAGCAGCGAGTCCCTACACACGCTGGCCGTCTTTGGCTCCATCATCATCTCCCTGTCCGTGTTTATGCTGATCGTGGCTAGCTTATGTTTCACCTACAAGCGACGCGAGAAGCTGCGCAAGCAGCCACGTCTCATCAACTCAATGTGCAACTCACAGCTGTCGCCTTTGTCACAACTGGTGGAACAGAGTTCGGGCGCCGGATCGGGATTACCATTGCTGGTGCAAAGAACCATTGCCAAGCAGATTCAGATGGTGCGACTGGTGGGCAAAGGACGATATGGCGAGGTCTGGCTGGCCAAATGGCGCGATGAGCGGGTGGCCGTCAAGACCTTCTTTACGACCGAAGAGGCTTCTTGGTTCCGCGAGACTGAAATCTATCAGACAGTGCTGATGCGACACGACAATATCTTGGGCTTCATTGCCGCCGATATCAAGGGTAATGGTAGCTGGACACAGATGTTGCTGATCACCGACTACCACGAGATGGGCAGCCTACACGATTACCTCTCAATGTCGGTGATCAATCCGCAGAAGCTGCAATTGCTGGCGTTTTCGCTGGCCTCCGGATTGGCCCACCTGCACGACGAGATTTTCGGAACCCCTGGCAAACCAGCTATCGCTCATCGCGATATCAAGAGCAAGAACATTTTGGTCAAGCGGAATGGGCAGTGCGCTATTGCTGACTTCGGGCTGGCAGTGAAGTACAACTCGGAACTGGATGTCATTCACATTGCACAGAATCCACGTGTCGGCACTCGACGCTACATGGCTCCAGAAGTATTGAGTCAGCAGCTGGATCCCAAGCAGTTTGAAGAGTTCAAACGGGCTGATATGTATTCAGTGGGTCTCGTTCTGTGGGAGATGACCCGTCGCTGCTACACACCCGTATCGGGCACCAAGACGACCACCTGCGAGGACTACGCCCTGCCCTATCACGATGTGGTGCCCTCGGATCCCACGTTCGAGGACATGCACGCTGTTGTGTGCGTAAAGGGTTTCCGGCCGCCGATACCATCACGCTGGCAGGAGGATGATGTACTCGCCACCGTATCCAAGATCATGCAGGAGTGCTGGCACCCGAATCCCACCGTTCGGCTGACTGCCCTGCGCGTAAAG AAGACGCTGGGGCGACTGGAAACAGACTGTCTAATCGATGTGCCCATTAAGATTGTC<u>AGATC</u> <u>T</u>CA-3’

Synthesized fragments were annealed and digested by NotI and BglII. Resultant two inserts (TkvS238A and GFP) were ligated to modified pPGW vector using NotI and AscI sites in the multiple cloning site. Transgenic flies were generated using strain attP2 by PhiC31 integrase-mediated transgenesis (BestGene Inc.).

### Generation of nos-loxP-mCherry-loxP-gal4-VP16 transgenic flies

Step 1: Construction of loxP-mCherry-SV40-loxP. mCherry cDNA was amplified using primers NheI mCherry Fw (5’-acgctagctatggtgagcaagggcgaggag-3’) and XhoI mCherry Rv (5’-gactcgagttacttgtacagctcgtccat-3’) from mCherry Vector (Clontech), and then the product was introduced into NheI-XhoI sites of the pFRT-SV40-FRT vector (Gift from Elizabeth R. Gavis). Step 2: BamHI-loxP-NotI oligo (5’-GATCCATAACTTCGTATAGCATACATTATACGAAGTTATGC-3’, 5’-GGCCGCATAACTTCGTATAATGTATGCTATACGAAGTTATG-3’) was inserted into BamHI NotI site after the SV40 polyA sequence of StepI vector. NdeI-loxP-NheI oligo (5’-CATATGCAACATGATAACTTCGTATAGCATACATTATACGAAGTTATTG-3’, 5’-CTAGCAATAACTTCGTATAATGTATGCTATACGAAGTTATCATGTTGCATATGCATG-3’) was inserted into the NdeI/NheI site upstream of mCherry sequence of Step 1 vector. Step 3: The NotI-BamHI flanked 3.13-Kb fragment from the pCSpnosFGVP (Gift from Elizabeth R. Gavis) containing the Nanos 5’ region-ATG (NdeI-start codon) Gal4-VP16-Nanos 3’ region was subcloned into NotI-BamHI sites of pUAST-attB. Step 4: The NdeI-flanked loxP-mCherry-SV40 polyA-loxP fragment was subcloned into the NdeI start codon of the plasmid described in Step 3. A transgene was introduced into the attP2 using PhiC31 integrase-mediated transgenesis systems by BestGene, Inc.

### Immunofluorescent Staining

Immunofluorescent staining was performed as described previously^34^. Briefly, testes were dissected in phosphate-buffered saline (PBS) and fixed in 4% formaldehyde in PBS for 30–60 minutes. Next, testes were washed in PBST (PBS + 0.3% TritonX-100) for at least 30 minutes, followed by incubation with primary antibody in 3% bovine serum albumin (BSA) in PBST at 4 °C overnight. Samples were washed for 60 minutes (three times for 20 minutes each) in PBST, incubated with secondary antibody in 3% BSA in PBST at 4 °C overnight, and then washed for 60 minutes (three times for 20 minutes each) in PBST. Samples were then mounted using VECTASHIELD with 4′,6-diamidino-2-phenylindole (DAPI) (Vector Lab, H-1200).

The primary antibodies used were as follows: 1B1, rat anti-Vasa (1:20) and mouse anti-Bam (1:20) were obtained from the Developmental Studies Hybridoma Bank (DSHB); Rabbit anti-Smad3 (phospho S423 + S425) antibody (1:100, Abcam, ab52903) AlexaFluor-conjugated secondary antibodies were used at a dilution of 1:400.

Images were taken using a Zeiss LSM800 confocal microscope with a 63× oil immersion objective (NA=1.4) and processed using Image J and Adobe Photoshop software. Three-dimensional rendering was performed by Imaris software.

### *In situ* hybridization

*In situ* hybridization on adult testes was performed as described previously^35^. Briefly, testes were dissected in 1X PBS and then fixed in 4% formaldehyde/PBS for 45 min. After fixing, they were rinsed 2 times with 1X PBS, then resuspended in 70% EtOH, and left overnight at 4°C. The next day, testes were washed briefly in the wash buffer (2X SSC and 10% deionized formamide), then incubated overnight at 37°C in the dark with 50 nM of Quasar® 570 labeled Stellaris probe against dpp mRNA (LGC Biosearch Technologies, kind gift from Michael Buszczak, previously validated^36^) in the Hybridization Buffer containing 2X SSC, 10% Dextran sulfate (MilliporeSigma), 1 μg/μl of yeast tRNA (MilliporeSigma), 2 mM Vanadyl ribonucleoside complex (NEB), 0.02% RNAse-free BSA (ThermoFisher), and 10% of deionized formamide. On the 3^rd^ day, testes were washed 2 times for 30 min each at 37°C in the dark in the prewarmed wash buffer (2X SSC, 10% Formamide) and then resuspended in a drop of VECTASHIELD with DAPI (Vector Lab, H-1200).

### Induction of Tkv-mCherry expression

For Tkv-mCherry clonal expression, hs-cre, nos-loxP-stop-loxP-Gal4 with UAS-Tkv-mCherry, UAS-GFP-αTubulin flies were heat-shocked at 37°C. for 15 min. Testes were dissected 24 hours after the heat shock. For the time course of Tkv-mCherry localization, hs-flp, nos-FRT-stop-FRT-Gal4 with UAS-Tkv-mCherry, UAS-GFP flies were heat-shocked at 37°C. for 60 min. Testes were dissected indicated times (day1, 2, 3) after the heat shock.

### Chloroquine or Lysotracker/LysoSensor treatment

Testes from newly eclosed flies were dissected into Schneider’s *Drosophila* medium containing 10% fetal bovine serum and glutamine–penicillin–streptomycin. Then testes were incubated at room temperature with or without 100 μM chloroquine (Sigma) in 1 mL media for 4 hours prior to imaging. For the lysosome staining, testes were incubated with 50 nM of LysoTracker Deep Red (ThermoFisher L12492) or 100 nM of LysoSensor Green DND-189 (ThermoFisher L7535) probes in 1 mL media for 10 min at room temperature then briefly rinsed with 1 mL of media for 3 times prior to imaging. These testes were placed onto Gold Seal™ Rite-On™ Micro Slides two etched rings with media, then covered with coverslips. An inverted Zeiss LSM800 confocal microscope with a 63× oil immersion objective (NA=1.4) was used for imaging.

### Quantification of pMad intensities

Mean intensity values in the GSC nuclear region were measured for anti-pMad staining. To normalize the staining condition, data were further normalized by the average of measurements of pMad from randomly picked three cyst cells in the same testes, and the ratios of relative intensities were calculated for each GSC (see Figure 3C).

### FRAP analysis

Fluorescence recovery after photo-bleaching (FRAP) of Dpp-mCherry signal was undertaken using a Zeiss LSM800 confocal laser scanning microscope with 63X/1.4 NA oil objective. Zen software was used for programming of each experiment. Encircled (approximately 10 μm diameter) areas of interest were photobleached using laser powers to achieve an approximately 70-90% bleach using the 561 nm laser. Fluorescence recovery was monitored every second.

### Statistical analysis and graphing

No statistical methods were used to predetermine sample size. The experiments were not randomized. The investigators were not blinded to allocation during experiments and outcome assessment. Statistical analysis and graphing were performed using Microsoft Excel 2010 or GraphPad Prism 7 software. Data are means and standard deviations. The P values (two-tailed Student’s t-test or adjusted *P* values from Dunnett’s multiple comparisons test) are provided.

---

**Figure S1.**
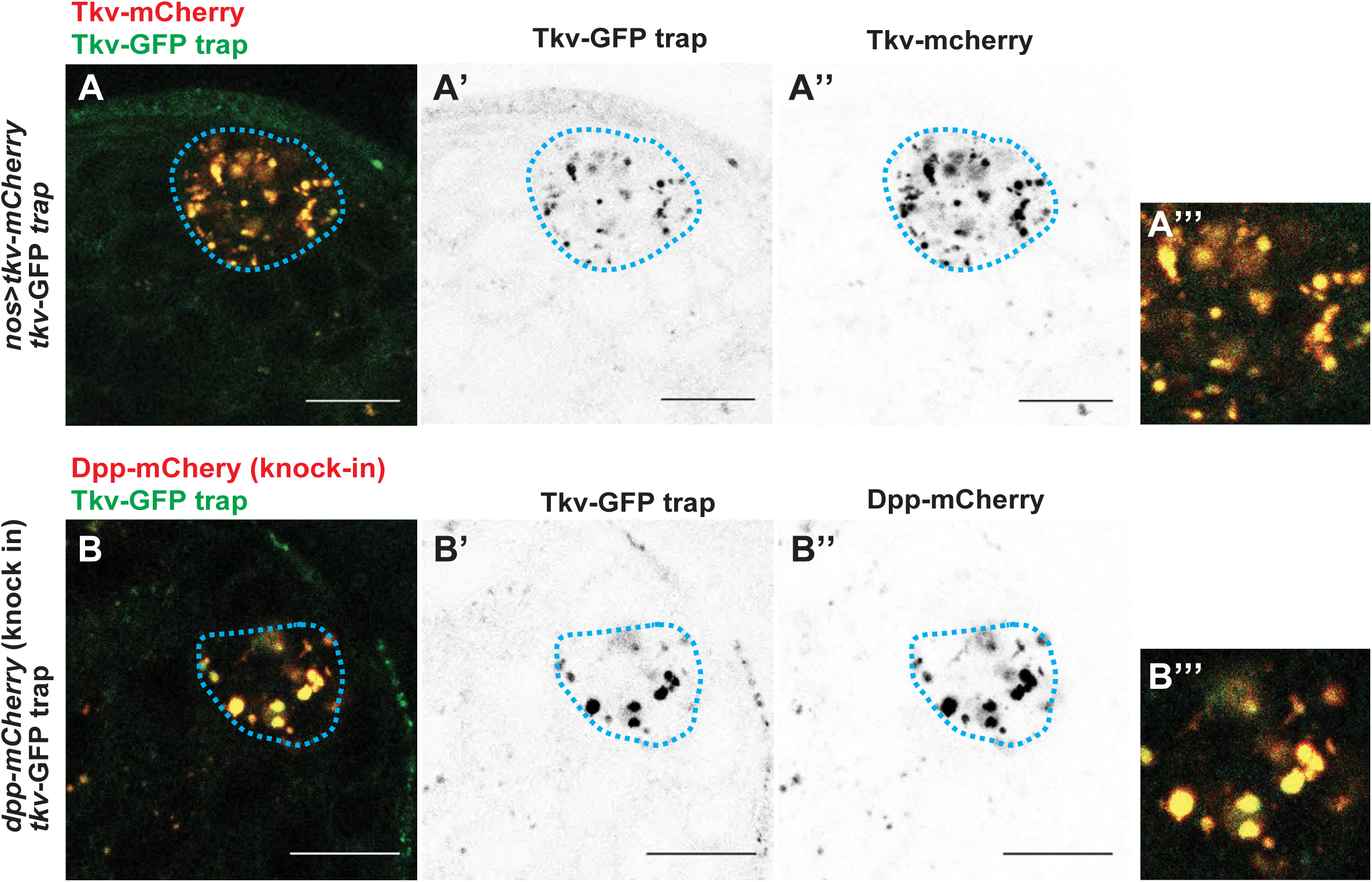
**(Associated to Figure1)** Representative images of the testis tip of indicated genotypes. **A**, Tkv-mCherry expressed in germline colocalizing with Tkv-GFP trap signal in the hub. **B**, Dpp-mCherry (CRISPR knock-in) signal colocalizing with Tkv-GFP trap signal in the hub. Scale bars, 10 μm. Blue dotted lines outline hub-GSC boundaries. Insets (**A’’’** and **B’’’**) show magnified hub area.

**Figure S2.**
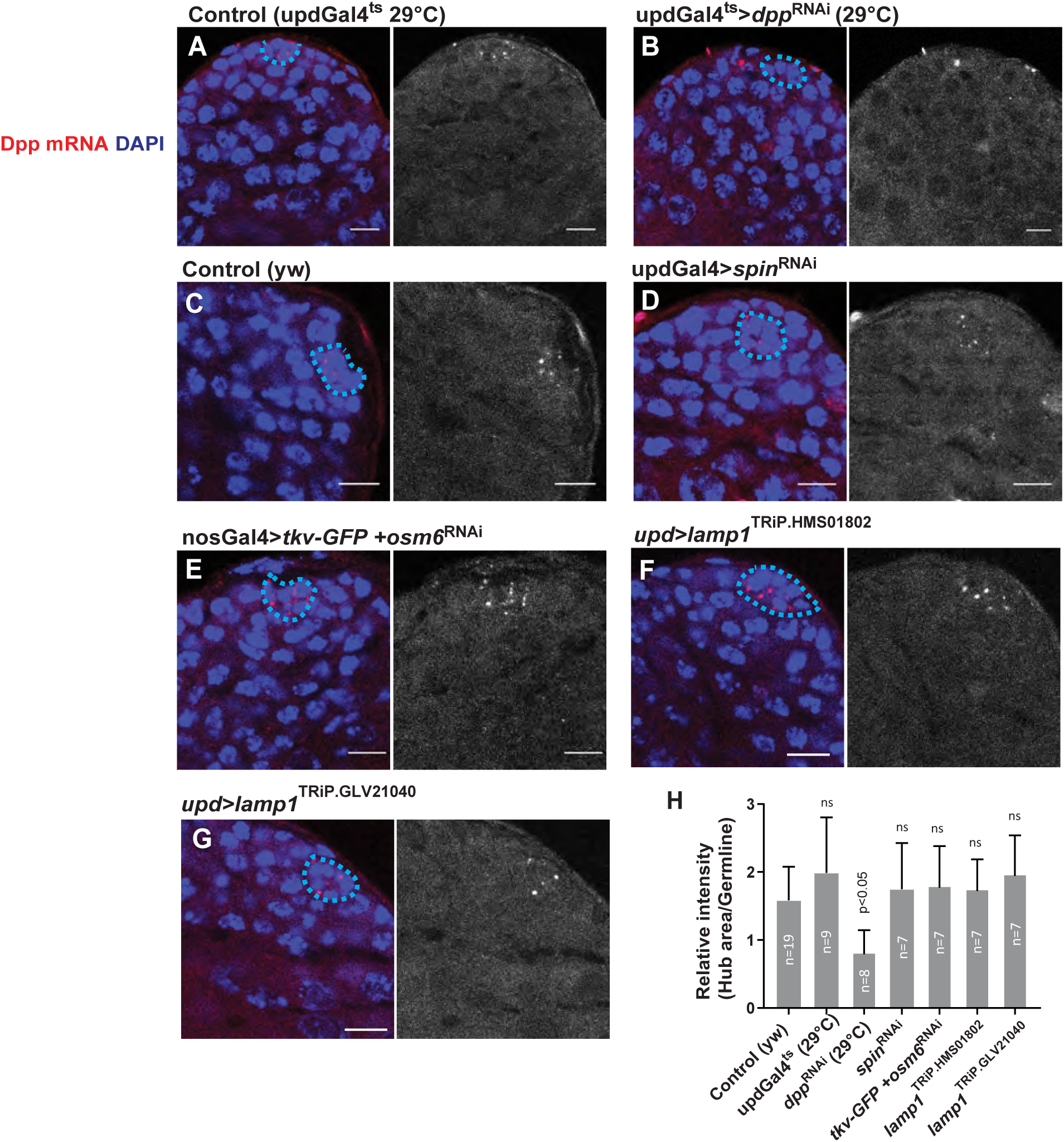
**(Associated to Figure2) A-G**, Representative images of *in situ* hybridization using Stellaris FISH probe against *dpp* mRNA (red) in the testis of indicated genotypes. **B**, Dpp RNAi (negative control) testis shows almost no detectable signal in the hub, suggesting the specificity of the probe. Hub is encircled by a blue dotted line. DAPI (blue) marks nuclei. Scale bars, 10 μm. **H**, Relative intensity values of Dpp in situ signal from indicated genotypes. Mean intensity values of entire hub area were divided by the average intensity from 2 randomly selected germline areas (5μm diameter circles) within same testis. Largest diameter of hub area was selected after the imaging of 0.5 μm z interval. The indicated numbers (n) of testes from two independent experiments were scored for each group. non-significant if not shown (P>0.05). Data are means and standard deviations. The adjusted *P* values from Dunnett’s multiple comparisons test are provided.

**Figure S3.**
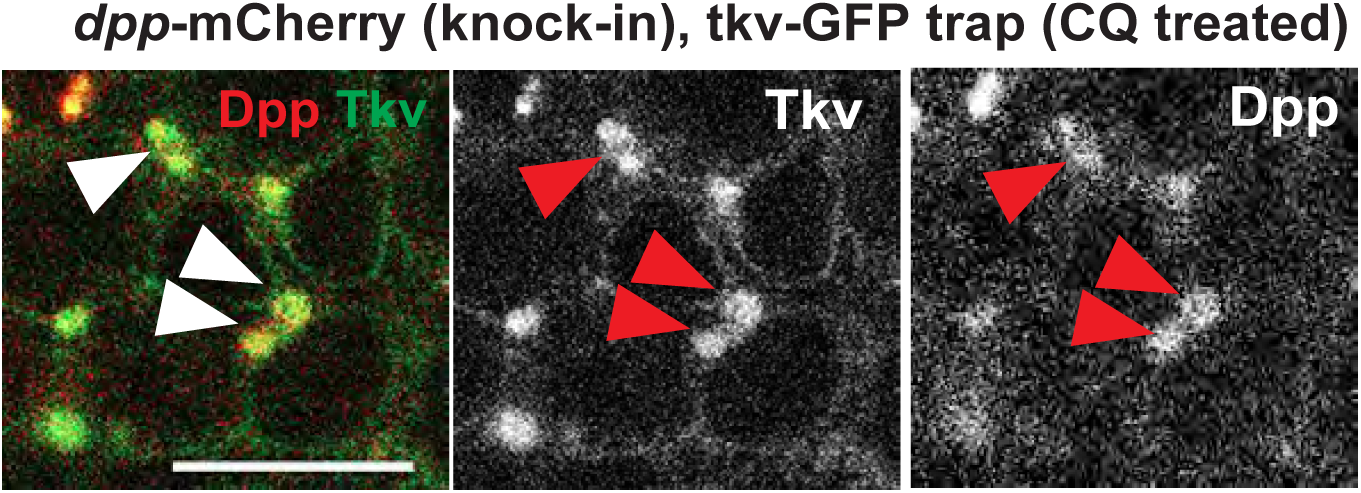
**(Associated to Figure4)** A representative image of the SG of Dpp-mCherry knock-in with tkv-GFP trap fly after 4-hour incubation with CQ. Arrowheads indicate enlarged lysosomes containing Dpp and Tkv. Scale b**4**ar, 10 μm.

## Supplementary movies (Associated to Figure4)

**Movie 1;** A representative video of quick recovery of the Dpp-mCherry signal after photo bleaching of the region away from the hub (Figure4I). Images were taken every second for 50 seconds.

**Movie 2;** A representative video of the Dpp-mCherry signal after photo bleaching of the entire hub region from the hub (Figure4J). Images were taken every second for 50 seconds.

